# Tandem: a bioinformatics tool for detection, mechanism classification, and population quantification of bacterial tandem gene duplications

**DOI:** 10.64898/2026.05.22.727201

**Authors:** Wing Yui Ngan, Ewan St. John Smith

## Abstract

**Motivation:** Tandem gene duplication drives antibiotic resistance, metabolic adaptation, and gene-family expansion in bacteria, but no tool detects them in reference genomes, discovers their junctions in isolate sequencing, and quantifies the junctions in population samples. Existing callers (e.g. *breseq*) detect duplications without classifying formation mechanisms and often fail to quantify the duplication.

**Results:** Tandem has 3 modules. Module 1 detects reference-genome duplications by NUCmer self-alignment and classifies each by homologous-recombination signature and the junction microhomology length. Module 2 confirms junctions in whole-genome sequencing at user-nominated coordinates after user inspecting the coverage plot. Module 3 quantifies known junction in population sequencing using the novel Junction Read Ratio (JRR). On 280 artificial population tests across seven bacterial species, Tandem achieves 100% recall and 4.3% mean absolute error. Applied to experimentally evolved *Pseudomonas fluorescens* SBW25 populations, Tandem resolves multiple co-segregating duplication fragments.

**Availability:** Source code, documentation, and test data are available under the MIT License at https://github.com/yuingan/tandem. Implemented in Python 3. Requires NUCmer (MUMmer4), minimap2, and samtools.

## 1. Introduction

Tandem gene duplications, adjacent copies of a genomic segment, are among the most common and consequential structural variants in bacterial genomes (Andersson and Hughes, 2009). They underlie gene amplification in antibiotic resistance (Sandegren and Andersson, 2009), metabolic adaptation in experimental evolution (Khomarbaghi, et al., 2024), and the expanding of virulence factor families (Maddamsetti, et al., 2024). Unlike point mutations, tandem duplications are inherently unstable. Copies can be lost through intramolecular recombination, making them a reversible and dynamic in genetic variation (Andersson and Hughes, 2009).

Each duplications could have different stability (Khomarbaghi, et al., 2024). The molecular mechanism by which a tandem duplication forms has direct consequences for its stability and evolutionary fate. Duplications mediated by homologous recombination (HR) between flanking direct repeats can lose the extra copy through the same repeats that created the duplication. In contrast, duplications formed by non-homologous end-joining (NHEJ), microhomology-mediated end-joining (MMEJ) or other non-HR repairing (e.g., alternative end-joining; Alt-EJ, single-strand annealing; SSA, synthesis-dependent MMEJ; SD-MMEJ, illegitimate recombination, etc) lack extended flanking homology and are therefore more stable. Experimentally, the mechanistic distinctions have been demonstrated in *Salmonella typhimurium* and *Pseudomonas fluorescens* SBW25 (Anderson and Roth, 1978; Ayan, et al., 2020; Khomarbaghi, et al., 2024) but has not been systematically assessed at genomic scale.

Existing bioinformatics tools address parts of this problem, but none provides an integrated solution. For example, *breseq* (Deatherage and Barrick, 2014) detects duplications in experimental evolution through new junction evidence, but it struggles with duplications near repetitive regions, does not classify the mechanism that generated them, and cannot quantify most duplications in population-level sequencing. A further tool is SegMantX (Hanke and Dagan, 2025), which detects diverged segmental duplications in bacterial plasmids but targets ancient duplications rather than young tandems. Lastly, general-purpose structural variant callers (DELLY, Manta, LUMPY) (Chen, et al., 2016; Layer, et al., 2014; Rausch, et al., 2012) are designed for diploid eukaryotic genomes and are poorly suited to haploid bacterial samples with population-level heterogeneity.

Here we present Tandem (Tandem Amplification and Duplication Event Mapper), a tool built for bacterial tandem duplications that integrates three analysis modules: (i) reference genome detection with mechanism classification, (ii) junction confirmation in sequences, and (iii) quantification of duplications. Tandem classifies each duplication by two independent features: the presence of an HR signature at outer boundaries (flanking repeat alignment) and the microhomology length at the inner junction. For non-HR duplications, Tandem avoids over-claiming pathway identity (e.g. NHEJ, MMEJ or Alt-EJ) and instead reports microhomology lengths for mechanism inference by users.

## 2. Materials and Methods

### 2.1 Overview

Tandem is written in Python 3 and consists of three modules, each addressing a distinct biological question (Fig 1). All modules use 1-based inclusive coordinates throughout, matching the NCBI/GenBank/IGV format. Tandem requires NUCmer (MUMmer4) for self-alignment, minimap2 for read mapping, and samtools for coverage analysis (Li, 2018; Li, et al., 2009; Marcais, et al., 2018). The tool can be installed via pip and provides a single command-line interface.

**Fig 1.**
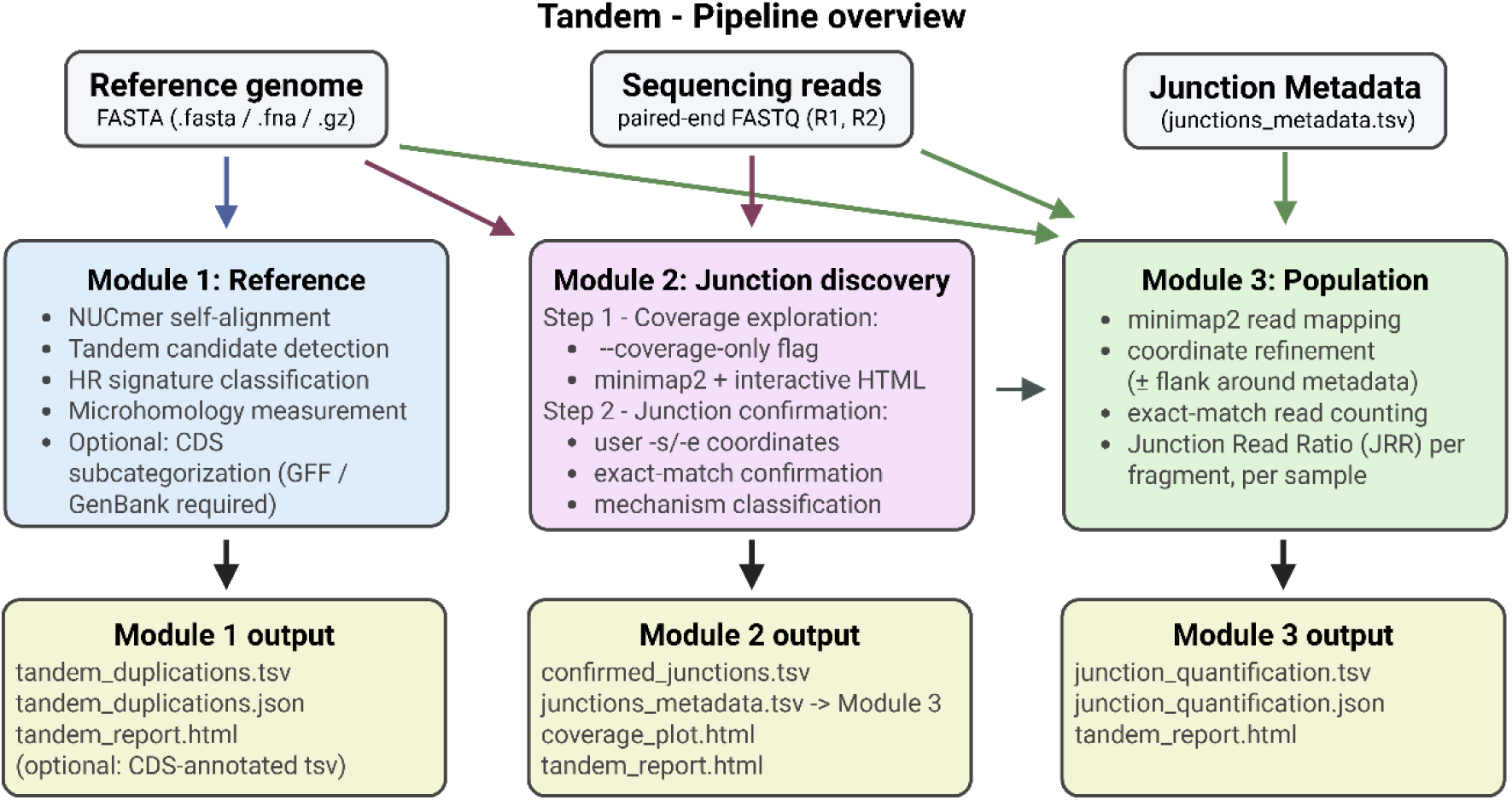
Tandem pipeline overview. Tandem has three modules sharing a common reference genome (top input). Module 1 detects tandem duplications in the reference genome by NUCmer self-alignment and classifies each duplication by HR signature (four scenarios for sequences mapped to reference with only one copy present; four checks for reference genome with two copies present), microhomology length, and optionally coding sequence (CDS) architecture. Module 2 confirms junctions in sequencing data in two steps. First, after coverage exploration (*--coverage-only*), users inspect the coverage and nominate candidate boundaries. Next, Module 2 confirms exact-match junction at user-supplied coordinates (*-iso -s*/*-e*). Module 2 generates junctions_metadata.tsv that can be used directly in Module 3. Users can also create the metadata file for junction quantification in Module 3. Module 3 quantifies known junction(s) in population sequences using the Junction Read Ratio (JRR), with ±20 bp coordinate refinement around metadata coordinates. Module 2’s junctions_metadata.tsv feeds directly into Module 3.

### 2.2 Module 1: Reference genome tandem detection

Module 1 detects tandem duplications in a reference genome through a three-stage pipeline:

#### Stage 1: Detection from sequence-geometric

NUCmer self-alignment identifies pairs of similar regions within the genome. Alignments are filtered by minimum identity (default 80%), minimum size (default 200 bp), and maximum inter-copy distance (default 30,000 bp). For circular genomes, distance is measured as the minimum of linear and wrap-around distance through the origin. Short-duplication filters (duplications < 500 bp) require the between copies to be shorter than the copy length and the copy sequence to have adequate k-mer complexity (at least 20% of possible 4-mers present), excluding mono/dinucleotide-repeat matches from replication slippage. Alternatively, coordinates from external tools (e.g. SegMantX, BISER, *breseq*) (Deatherage and Barrick, 2014; Hanke and Dagan, 2025; Iseric, et al., 2022) can be provided via *--detection-input*, bypassing NUCmer.

#### Stage 2: CDS subcategorization (optional)

When a GFF3 or GenBank annotation is provided (*--annotation*), each duplication is classified by its CDS content: intergenic, tandem single-gene, tandem segmental, or proximal, based on the number of CDSs fully contained in each copy and the intervening gap. This improves user experience in interpretation of the duplicated regions for their biological discoveries/questions.

#### Stage 3: Mechanism classification

Each duplication is classified by two independent features. Tandem first detects the HR signature detection using the following procedures. For a tandem duplication with two NUCmer-reported copies, the tool searches for extended homology (direct repeats) at the outer boundaries. This addresses the intrinsic NUCmer bracketing ambiguity of an R-D-R-D-R structure (where R is the repeat and D is the duplication) by testing both bracketing scenarios and comparing both copies’ R region against the external R in each scenario (Fig 2A and B). Aligned windows are scored strictly (match +2, mismatch -3, gap open -5, gap extend -2) so that only near-identical stretches score positively, and a sliding-window identity analysis (high-scoring segment pair; HSP) finds the best sub-region of at least 35 bp at 92% or higher identity. A complexity filter rejects HR calls when both flanking windows are low-complexity (microsatellite periodicity), distinguishing replication slippage from true HR; this can be disabled with *--no-hr-complexity-filter*. Next, Tandem assesses the microhomology at the inner junction. The microhomology is measured as the maximum exact overlap between the end of copy 1 and the start of copy 2.

**Fig 2.**
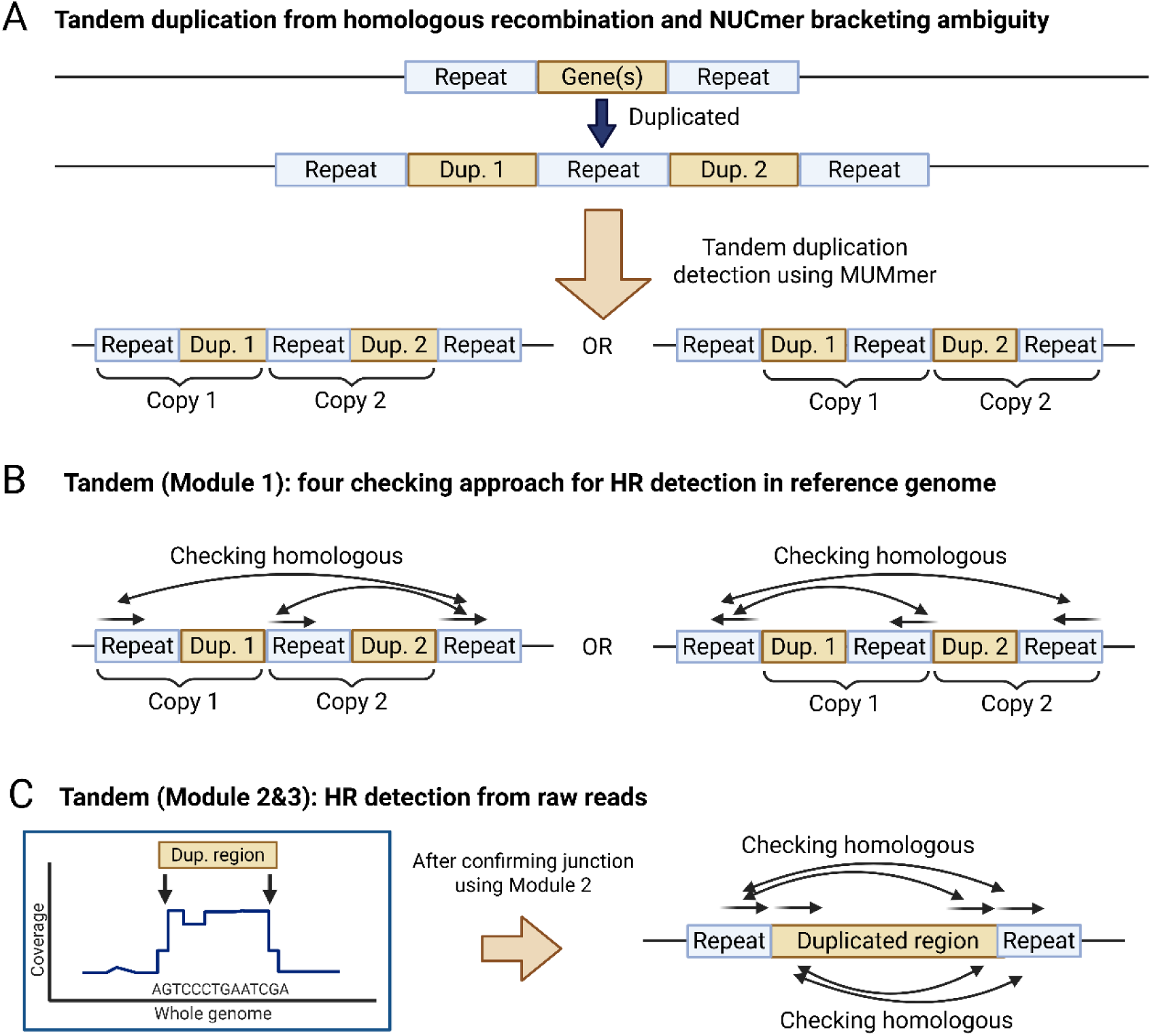
HR detection geometry in Tandem. **(A)** The R-D-R-D-R model for HR-mediated tandem duplication. NUCmer has an intrinsic bracketing ambiguity: the duplicated sequence can be reported as [R-D][R-D]-R (Scenario 1, R at the start of each copy) or R-[D-R][D-R] (Scenario 2, R at the end). **(B)** Module 1 tests both scenarios, aligning each copy’s repeat region against the external repeat using a sliding-window high-scoring segment pair (HSP) analysis with a complexity filter to exclude microsatellite periodicity. **(C)** Single-copy classifier used in Modules 2 and 3. Since only one copy of the duplicated region D exists in the reference, the tool searches for direct repeats flanking D under four mechanistically distinct scenarios with s1/s2 (asymmetric: outside on one side, inside at the boundary on the other), s3 (repeats outside D on both sides; classical R-D-R), and s4 (repeats inside D at both internal boundaries, mediating sister-chromatid unequal crossing-over). The outward search window scales adaptively with duplication size (up to 5 kb for ≥ 50 kb duplications) to capture flanking rRNA/IS elements. Microhomology at the junction is the longest exact match between the end of the duplicated region and its start.

Tandem reports mechanism-classification confidence based on duplication size, reflecting how reliably the flanking-repeat search can be performed rather than the biological mechanism itself. Because the outward HR search window scales with duplication size, very small duplications (< 300 bp) yield a search window too small to reliably detect or exclude a flanking repeat (low confidence). Duplications of 300-500 bp give moderate confidence and duplications ≥ 500 bp give a search window large enough for robust flanking-repeat inference (high confidence). This aims to provide extra information for the reliability of the HR/no-HR call, not the certainty that any particular mechanism produced the duplication.

### 2.3 Module 2: Junction discovery

Module 2 confirms novel duplication junctions from whole-genome sequencing data with known reference. The workflow consists of two steps and requires the user to interpret the candidate junction positions from the coverage plot. Firstly, with *--coverage-only* flag, Tandem maps sequencing reads to the reference with per-base coverage in sliding windows and generates an interactive HTML coverage plot. The user inspects this plot to nominate candidate duplicated regions, typically plateau-shaped coverage elevations with sharp boundaries. This step accepts both isolate and population sequences, although coverage-based detectability of low-frequency duplications is bounded by the population frequency (a duplication carried by 5% of cells produces only ∼5% coverage elevation, near the threshold of visual inspection). In the junction-confirmation step (*-iso* with user-supplied *-s*/*-e* coordinates), Tandem constructs a candidate junction sequence for each pair of nominated boundaries recreating the novel sequence that exists only when the two copies are placed in tandem. The junction sequence begins at 15 bp on each side of the boundary and is extended symmetrically, one base per side at a time, until it is unique in the reference genome (a junction must be unique before it can be quantified, since a non-unique probe would match elsewhere). The reads are then searched for each unique junction by exact match. Only reads containing the junction as a contiguous, byte-identical substring (no gaps, no mismatches) are counted. This junction generation is parallelized across user-supplied threads with dynamic load balancing, partitioning the candidate search space into ten times as many chunks as worker processes so that time-consuming candidates in repetitive regions do not block fast ones. Each confirmed junction is annotated with HR signature, microhomology length, and supporting read count using the single-copy classifier described in section 2.5.

### 2.4 Module 3: Population junction quantification

Module 3 quantifies the known duplication junctions in population sequencing data. Given approximate junction coordinates from metadata, the tool searches a small neighbourhood (default ± 20 bp) to find the exact junction position, mitigating coordinate uncertainty from microhomology or different reporting conventions. Junction reads are counted by exact match and the Junction Read Ratio (JRR) is calculated as:

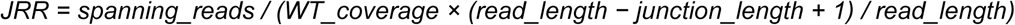

where WT_coverage is the genome-wide mean coverage from minimap2 mapping.

The JRR is biased low relative to true frequency because (i) exact-match counting excludes reads containing one or more sequencing errors, and (ii) the junction-uniqueness extension increases the length over which a read must perfectly match, reducing the number of qualifying read positions. This bias is assessed and well-characterised by the 4.3% mean absolute error reported in section 3.3. JRR should therefore be interpreted as a calibrated relative indicator suitable for tracking duplication dynamics across time-points and conditions, rather than as an absolute allele-frequency estimate. Mechanism classification uses the same single-copy classifier as Module 2.

### 2.5 Duplication mechanism identifier from sequencing data (Modules 2 and 3)

When a duplication exists in an isolate or population sample, only a single copy is present in the reference genome. The boundary geometry differs from Module 1’s two-copy case (Fig 2). In Module 1, both duplicated copies are visible directly in the reference, so the four-check approach (Fig 2A and 2B) sequentially tests four alignment pairings (R₁ vs R₃, R₂ vs R₃, R₂ vs R₁, and R₃ vs R₁), returning a positive HR call as soon as any one of them passes the alignment criteria.

This approach covers all positions where the original mediating homology could have sat. For Modules 2 and 3, the classifier examines homology around the confirmed start and end of the duplication, and the sequence at the inner junction (Fig 2C). Tandem’s built-in *classify_single_copy_junction* function therefore performs two independent operations against the reference. First, HR detection searches for direct repeats flanking the duplicated region D using four mechanistically distinct scenarios, each tested with the same windowed high-scoring segment pair (HSP) analysis (≥ 35 bp at ≥ 92% identity, with the complexity filter applied), and the longest valid match is reported. Scenario s3 (primary) tests for repeats lying strictly outside D on both sides with the classical R-D-R configuration in which two flanking direct repeats recombine intramolecularly to produce a tandem. Scenarios s1 and s2 are asymmetric: s1 tests an outside-flanking repeat at the start against an internal repeat at the end (the duplication boundary at the end cuts internally through a repeat copy); s2 is the mirror. Scenario s4 tests for repeats lying entirely inside D, at the immediate internal start and end of the duplicated region, capturing duplications produced by sister-chromatid unequal crossing-over between two internal homologous tracts (e.g. rRNA operons or IS elements positioned at the inner boundaries of D).

The four scenarios use strictly disjointed window halves so that scenario labels report the underlying mechanism unambiguously. Second, microhomology at the inner junction is measured as the longest exact match between the END of the duplicated region D and the START of D targeting the biological junction sequence created when copy 1’s end is joined directly to copy 2’s start. The HR signature provides the primary mechanism classification and the microhomology length provides additional sequence-level context useful for distinguishing among non-HR mechanisms (e.g. MMEJ at 5-25 bp microhomology, Ku/LigD-type NHEJ at 0-4 bp) when no HR signature is present.

To detect flanking repeats in large duplications, where rRNA operons or IS elements may sit several kb outside the boundary, the outward search window scales adaptively with duplication size (200 bp for duplications < 10 kb, 2 kb for 10-50 kb, and 5 kb for ≥ 50 kb). The same alignment scoring and complexity filter as Module 1 are applied.

An absent HR signature is a statement about substrate, not about RecA activity. RecA is the central bacterial recombinase that catalyses homology search and strand exchange during homologous recombination (Bell and Kowalczykowski, 2016; Cox, 2007). Classical RecA-mediated HR between flanking direct repeats generates tandem duplications with the boundary geometry Tandem detects. The biological interpretation of HR-negative duplications, including replication template switching and Ku/LigD-type non-homologous end-joining (NHEJ), is discussed in the Discussion section.

### 2.6 Comparison with existing tools

We compared Tandem against *breseq* v0.39 (Deatherage and Barrick, 2014) for isolate-level junction discovery on the same SBW25 experimental evolution dataset (Khomarbaghi, et al., 2024). *breseq* was run with default parameters using the SBW25 GenBank reference (NC_012660.1). Tandem Module 2 was run with default parameters. For reference-genome analysis, we provide a conceptual scope comparison between Tandem Module 1 and SegMantX (Hanke and Dagan, 2025). We did not run SegMantX in this work, since the two tools target different biological questions (recent tandem versus ancient diverged duplications).

### 2.7 Hardware and software environment

All benchmarks reported in this work were performed on a Dell Precision 7875 Tower workstation equipped with an AMD Ryzen Threadripper PRO 7965WX processor (24 physical cores, 48 threads) and 256 GB of system memory, running Ubuntu 24.04.3 LTS under Windows Subsystem for Linux 2 (WSL2; kernel 6.6.87.2-microsoft-standard-WSL2). Tandem was executed with 44 threads (*-t 44*) for all multi-threaded operations. External dependencies were NUCmer (MUMmer 4.0.1), minimap2 (2.30-r1287), samtools (1.23.1), and bowtie2 (2.5.5), installed within a conda environment running Python 3.12.13. Runtime measurements were captured with GNU *time -v*.

## 3. Results

### 3.1. Module 1: Tandem duplication detection across 8 bacterial genomes

We tested Tandem Module 1 on eight reference genomes selected across diversity and most relevant to tandem duplication biology. These considerations include genome size (0.58-8.67 Mb), GC content (32-72%), repeat content (from minimal *Mycoplasma* genomes to the repeat-rich PE/PPE architecture of *Mycobacterium tuberculosis*), DNA repair complement (NHEJ-capable and NHEJ-absent species), and phylogenetic diversity (Proteobacteria, Actinobacteria, and Mollicutes). Module 1 completed all eight reference genomes in under 40 seconds total (per-genome times ranging from 1.8 s for *Mycoplasma pneumoniae* to 9.9 s for the 6.72 Mb *Pseudomonas fluorescens* SBW25 chromosome; Table S3). *P. fluorescens* SBW25 was included specifically because it has published experimental evolution data with duplication boundaries (Khomarbaghi, et al., 2024), which support Module 2 validation in section 3.2. Across these eight genomes, Tandem detected between 4 and 53 tandem duplications per genome (Table 1), with *Mycobacterium tuberculosis* H37Rv showing the highest count consistent with its PE/PPE repeat-rich architecture. The fraction of duplications classified as carrying an HR signature ranged from 0% (*M. pneumoniae* M129) to 58.6% (*P. fluorescens* SBW25), consistent with SBW25’s repeat-rich genome and active recombination machinery.

**Table 1.**
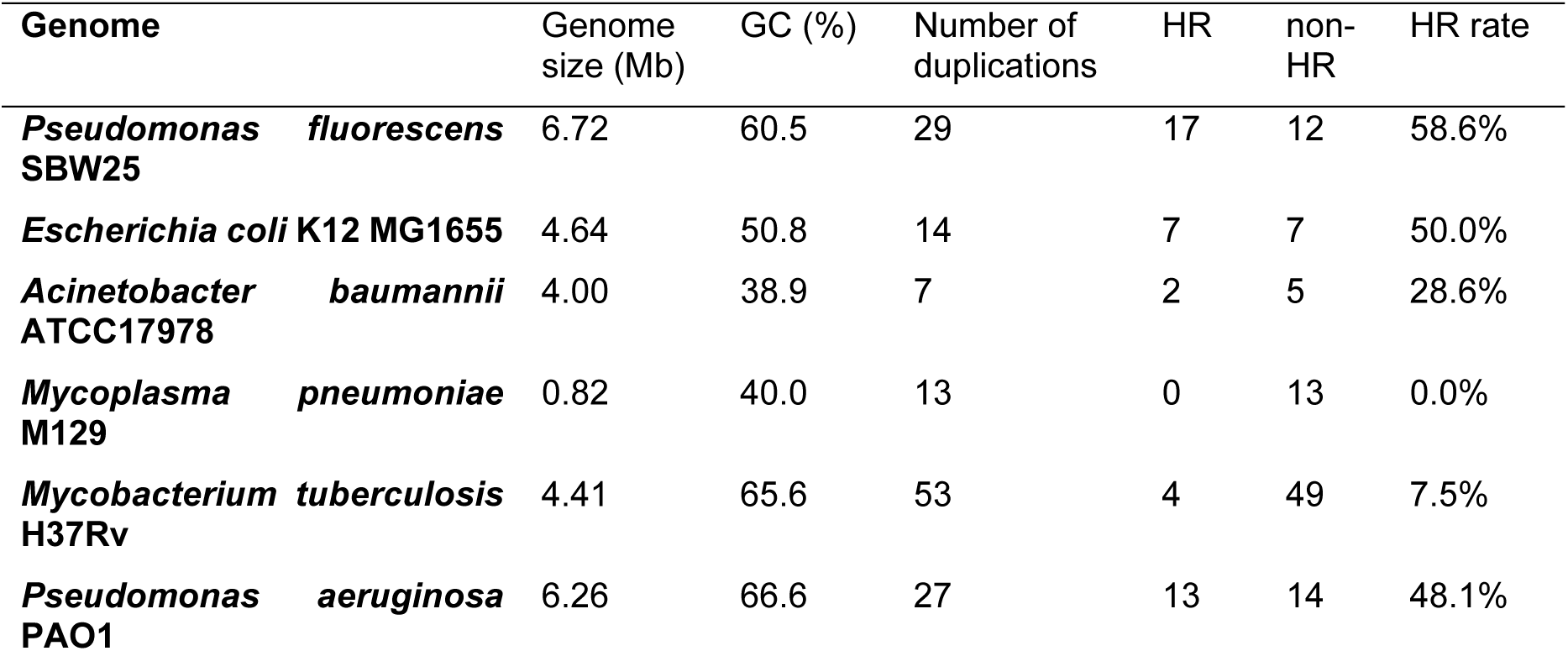
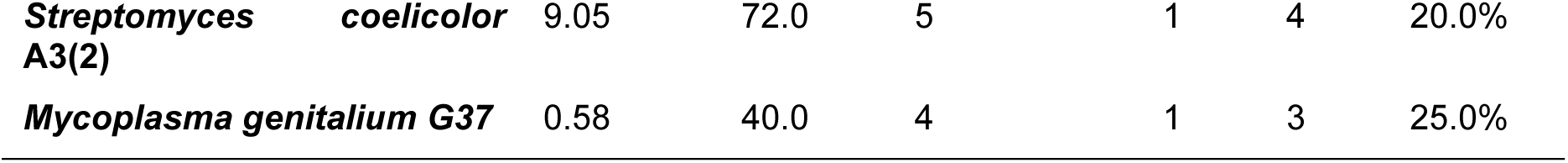
Module 1 detection across eight reference genomes.

HR sensitivity analysis (SBW25, Supplementary Table S1) confirmed that the HR classification is stable across reasonable parameter ranges. Default (≥35bp at ≥92% identity) and permissive (≥30bp at ≥90% identity) settings yielded near-identical results, with 17 and 18 HR-positive duplications out of 29 respectively, identical mean duplication size of 538 bp, and matching confidence-level distributions. Under strict settings (≥50bp consecutive match at ≥95% identity combined with stricter NUCmer detection parameters), only 11 high-confidence duplications were detected (mean size 931 bp), none of which met the strict HR threshold. This suggests most HR signatures in SBW25 fall in the 35 to 100 bp range at 92 to 96% identity, which is biologically reasonable for rhs-like repeat elements and intermediate IS fragments, and is not captured by the strict 50bp at 95% criterion. We adopted default thresholds as the primary settings because they correctly recovered the manually annotated HR/non-HR classifications in published SBW25 isolate data.

### 3.2 Module 2: Junction discovery in SBW25 experimental evolution

We applied Tandem Module 2 and *breseq* to the five sequenced isolates (M1-M5) from a *Pseudomonas fluorescens* SBW25 experimental evolution experiment (Khomarbaghi, et al., 2024) to detect the duplication junctions in each isolate. Tandem additionally reported mechanism classification (HR signature, microhomology length) for each junction (Table 2). M1’s single junction (1.02 Mb) lacked an HR signature and showed 3 bp microhomology at the junction, consistent with end-joining or replication-mediated formation rather than classical homology-driven amplification. Across M2-M5, Tandem identified an HR signature with asymmetric flanking-repeat geometry. M2’s nine HR-positive variants all carry a 428 bp match at 92.1% identity in scenario s1 (outside-flanking repeat at the start, internal repeat at the end), M3’s fourteen variants and M4’s two variants carry scenario s2 matches (internal repeat at the start, outside-flanking repeat at the end), with s2 match lengths of 136-442 bp (M3) and 1544-1772 bp (M4) and M5’s ten variants split between scenarios s1 (3 variants) and s2 (7 variants), reflecting alignment ambiguity at this boundary. The unified scenario calls per biological event in M2, M3, and M4 and the predominantly-s2 distribution in M5 are consistent with the published boundary coordination (Khomarbaghi, et al., 2024), which report *rhs* genes boundaries on one side and intergenic-or-no-repeat regions on the other for these duplications.

**Table 2.** Tandem vs *breseq* comparison on *P. fluorescens* SBW25 isolates M1-M5. Each isolate carries a single major tandem duplication. Tandem reports multiple junction-row variants per event reflecting microhomology-driven coordinate uncertainty, which we collapse here to one event per isolate (representative coordinates and total spanning reads shown). breseq detected 2/5 events as new-junction evidence (JC). 0/5 were detected as large duplications (LD) by *breseq*’s coverage-based caller.

| Sample | Dup. size | Tandem detection* | Tandem mechanism** | *breseq* detection | Concordance |
| --- | --- | --- | --- | --- | --- |
| **M1** | 1.02 Mb | 1 event, 1 variant<br>(21 reads) | no HR; MH = 3 bp | JC at<br>1848645/2864<br>810 (26 reads) | concordant |
| **M2** | 1.01 Mb | 1 event, 9 variants<br>(27 reads) | HR sig (s1, 428 bp<br>at 92.1% identity) | absent | Tandem only |
| **M3** | 490 kb | 1 event, 14 variants<br>(47 reads) | HR sig (s2, 136 bp<br>at 92.7% identity) | JC at<br>2296498/2786<br>206 (37 reads) | concordant |
| **M4** | 1.01 Mb | 1 event, 2 variants<br>(2 reads) | HR sig (s2, 1772<br>bp at 95.7%<br>identity) | absent | Tandem only |
| **M5** | 1.01 Mb | 1 event, 10 variants<br>(22 reads) | HR sig (s2 ×7 & s1<br>×3, 105-381 bp at<br>92.1-92.6%<br>identity) | absent | Tandem only |

\* Variants are due to microhomology at the junction makes the exact boundary ambiguous. The junction has higher confidence with more reads and Tandem does not filter junctions with low read counts allowing user to interpret raw results.
\*\*In M5, multiple sequences fall into multiple scenarios.

Tandem reports every confirmed junction position independently. Duplications from microhomology-rich boundary regions can produce multiple junction-position variants for the same biological event. The dominant variant typically accounts for the majority of reads, with the remainder supported by singletons that represent alignment-position uncertainty rather than independent events. For isolate analysis, users may filter variants by minimum read support to focus on dominant junctions. For population sequencing, where low-frequency subpopulations are biologically meaningful, no such filter should be applied, since singletons may represent rare but real subpopulations.

M4 has the lowest read support of any isolate, with only 2 spanning reads across 2 variants. This likely reflects the duplication boundaries falling within highly repetitive regions, where even extended junction sequences are not unique to the duplication (repeated elsewhere in the chromosome) and so cannot be matched by short reads. Despite the low read count, the HR match of 1544-1772 bp at over 95% identity is itself strong evidence for the call.

### 3.3 Module 3: Population-level quantification from Junction Read Ratio

We validated Module 3 using artificial populations generated by in-silico mixing of simulated mutant and wild-type reads at known frequencies across seven bacterial species previously assessed in Module 2, four time points each, and ten mutant junctions per species (280 tests total). Across all species, Module 3 detected every spiked junction (100% recall) with mean absolute error ranging from 3.6% (*P. aeruginosa*) to 5.2% (*E. coli*) and an aggregate MAE of 4.3% (Table 3, Figure 3).

**Fig 3.**
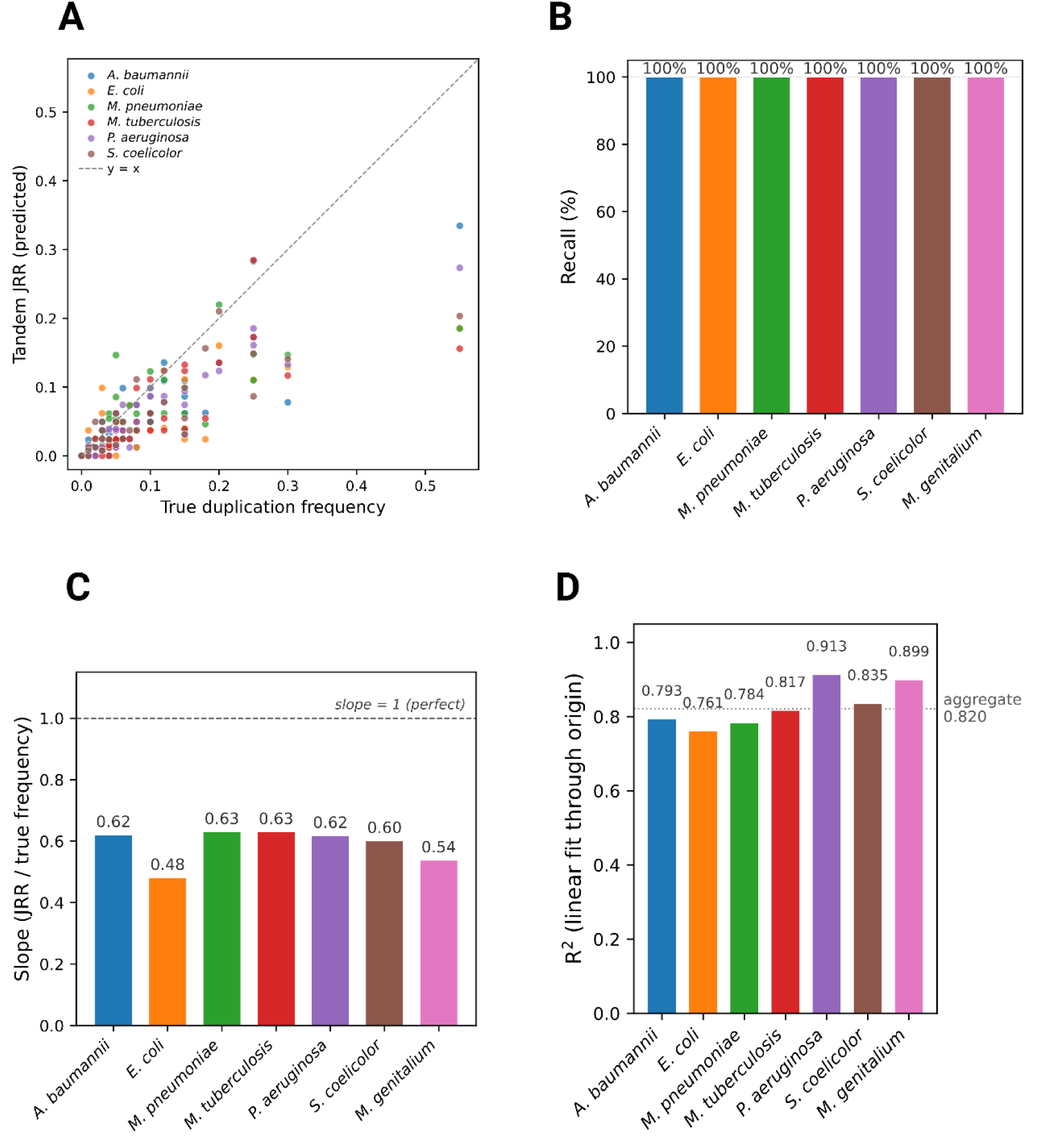
Module 3 validation on artificial populations across seven bacterial species. **(A)** Predicted JRR vs true duplication frequency for all 280 tests, color-coded by species. Dashed line is y = x. **(B)** 100% per-species recall across all seven species (n = 40 tests per species). **(C)** Per-species regression slope (JRR / true frequency) in the 0-30% linear range, fit through the origin. Dashed reference at slope = 1 indicates perfect calibration. The aggregate slope of ≈ 0.60 quantifies the systematic underestimation arising from exact-match counting and unique-junction extension (section 2.4), and is consistent across most species. **(D)** Per-species R² of the same through-origin linear fit; dotted reference is the aggregate R² of 0.81 across all species pooled. The high R² values demonstrate that JRR is a reliable relative indicator suitable for tracking duplication dynamics across time-points.

**Table 3.** Module 3 validation on artificial populations across seven bacterial species (n = 40 per species; 4 time points × 10 mutants).

| Species | n tests | Detected | Recall | MAE |
| --- | --- | --- | --- | --- |
| *A. baumannii* ATCC17978 | 40 | 40 | 100% | 0.0407 |
| *E. coli* K12 MG1655 | 40 | 40 | 100% | 0.0518 |
| *M. pneumoniae* M129 | 40 | 40 | 100% | 0.0417 |
| *M. tuberculosis* H37Rv | 40 | 40 | 100% | 0.0452 |
| *P. aeruginosa* PAO1 | 40 | 40 | 100% | 0.0359 |
| *S. coelicolor* A3(2) | 40 | 40 | 100% | 0.0413 |
| *M. genitalium* G37 | 40 | 40 | 100% | 0.0443 |
| Total/aggregate | 280 | 280 | 100% | 0.0430 (4.3%) |

Application to experimentally evolved *P. fluorescens* SBW25 populations at day 28 successfully quantified multiple co-segregating duplication fragments (Table 4, Figure 4).

**Figure 4.**
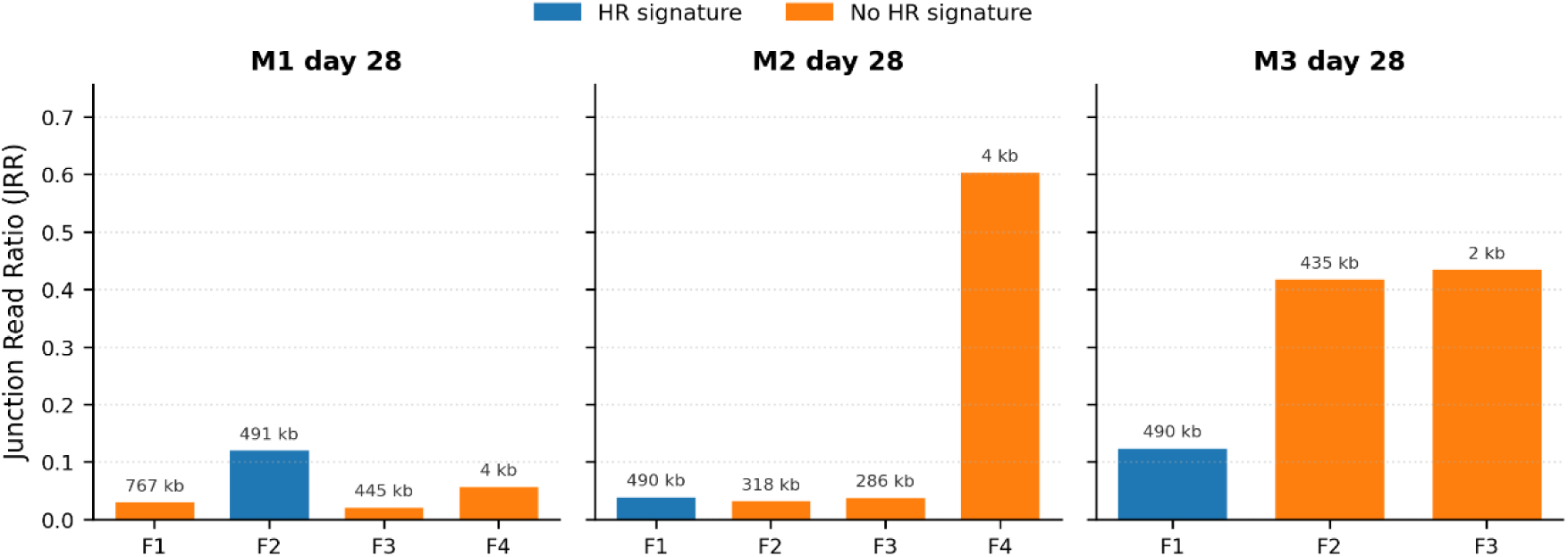
JRR per fragment in three *P. fluorescens* SBW25 day-28 populations. M2 is dominated by the small (3.6 kb) F4 fragment at JRR = 0.61. M3 shows two large fragments at comparable abundance (F2 at 435 kb, F3 at 2.2 kb; JRR ≈ 0.42-0.44). M1 has a single dominant HR-positive fragment (F2, 491 kb, JRR = 0.12). Bar size labels indicate duplication length; HR/no-HR coloring are scenario classification results from Tandem.

**Table 4.** *P. fluorescens* SBW25 M1, M2, and M3 population quantification at day 28 (Module 3, day-28 metadata).

| Population | Fragment | Coordinates | Size | JRR | HR signature | MH (bp) |
| --- | --- | --- | --- | --- | --- | --- |
| M1 | F1 | 2,375,280-3,142,292 | 767 kb | 0.032 | no | 0 |
| M1 | F2 | 2,296,304-2,786,885 | 490.6 kb | 0.121 | yes | 2 |
| M1 | F3 | 2,267,938-2,712,554 | 444.6 kb | 0.022 | no | 0 |
| M1 | F4 | 2,375,695-2,379,967 | 4.3 kb | 0.058 | no | 0 |
| M2 | F1 | 2,296,925-2,787,139 | 490.2 kb | 0.041 | yes | 0 |
| M2 | F2 | 2,347,702-2,666,045 | 318.3 kb | 0.034 | no | 0 |
| M2 | F3 | 2,239,941-2,525,738 | 285.8 kb | 0.039 | no | 1 |
| M2 | F4 | 2,375,566-2,379,196 | 3.6 kb | 0.605 | no | 1 |
| M3 | F1 | 2,296,498-2,786,206 | 489.7 kb | 0.125 | yes | 0 |
| M3 | F2 | 2,374,401-2,809,371 | 435.0 kb | 0.420 | no | 0 |
| M3 | F3 | 2,374,589-2,376,758 | 2.2 kb | 0.436 | no | 1 |

In the M2 population, the F4 fragment (3.6 kb) dominated at JRR = 0.61, consistent with a small high-frequency tandem amplification. In M3, the F3 (2.2 kb) and F2 (435 kb) fragments co-existed at comparable frequencies (JRR = 0.44 and 0.42 respectively). For the M3-F1 fragment (489.7 kb), which identified an HR signature (asymmetric flanking-repeat geometry, scenario s2, 136 bp at 92.7% identity), was present at a lower frequency (JRR = 0.13). In M1, the only fragment with non-trivial frequency was F2 (490.6 kb, HR-positive, JRR = 0.12). These frequency estimates by JRR are consistent with the experimental evolution dynamics reported in (Khomarbaghi, et al., 2024).

Notably, several large duplications (285-767 kb across M1-M3 populations) carry no HR signature and exhibit MH = 0-1 bp at the junction. The absence of detectable flanking direct repeats and the very short junction microhomology argue against both classical RecA-mediated HR and against MMEJ (which typically operates with 5-25 bp microhomology). Possible alternatives include replication template switching at stalled forks and Ku/LigD-type NHEJ. This is further discussed in the discussion.

### 3.4 Comparison with *breseq*

We ran *breseq* v0.39.0 on the same five *P. fluorescens* SBW25 isolates with default parameters and the *P. fluorescens* SBW25 GenBank reference (NC_012660.1). *breseq* reports structural variants with two evidence categories. Large-duplication (LD) entries based on elevated read-depth, and new-junction (JC) entries based on split-read evidence. Across all five isolates, *breseq* emitted zero LD entries; structural-variant detection therefore depended entirely on JC evidence. At the biological-event level, *breseq* detected the duplication junction in 2/5 isolates (M1 and M3) with concordant coordinates (within 200 bp of Tandem’s representative junction) and similar read support (26 and 37 reads respectively). *breseq* did not detect any junction near the duplication boundaries in M2, M4, or M5. The relevant regions (∼1853 kb / ∼2865 kb) had no JC entries at all in those samples, neither passing nor failing *breseq*’s quality filters.

The discrepancy is attributable to microhomology-driven dispersion of the junction read alignments. In M2, M4, and M5 the duplication junction sits inside a region of short microhomology (0-3 bp), and the resulting reads align ambiguously across a ∼5 kb window of nearly equivalent breakpoint coordinates. Tandem reports each distinct junction sequence as a separate row but recognises them as variants of the same biological event when they share duplication size (*dup_size*) and approximate boundaries. In these three isolates the spanning reads are spread across 9, 2, and 10 coordinate variants respectively (Table 2). *breseq*’s JC caller appears to require a single coordinate to accumulate enough reads to pass its threshold, and the microhomology-induced spread leaves no individual coordinate above that threshold. M1 (where Tandem reports a single junction with no microhomology spread) and M3 (where 31 of 47 reads concentrate at two adjacent coordinates) are the cases where *breseq*’s threshold is satisfied.

Beyond raw detection, Tandem provides three pieces of information that *breseq* does not. First, in mechanism classification, Tandem labels each junction with HR signature status (4/5 isolates HR-positive, with asymmetric flanking-repeat geometry. Tandem identified scenario s1 in M2, scenario s2 in M3 and M4, mixed s1/s2 in M5) and microhomology length (0-3 bp), supporting biological inference that *breseq*’s output cannot. Second, for junction clustering, Tandem’s recognition that 9 junction-row variants in M2 represent the same biological event collapses what *breseq* sees as scattered low-support evidence into a single high-confidence call. Third, for population-level quantification, Tandem’s Module 3 has no *breseq* comparator. *breseq* failed to report most of the duplications because duplication frequency cannot be easily measured using the same approach to measure Single Nucleotide Polymorphisms (SNP) frequency. To overcome this difficulty, Tandem has largely advanced the quantification method by introducing JRR. *breseq* runtimes ranged from 38.5 to 42.4 minutes per isolate (44 threads), comparable to Tandem Module 2’s 31-58 minutes per isolate on the same hardware. Both tools are sufficiently fast for routine use in bacterial experimental evolution pipelines and can run on most laptops.

### 3.5 Conceptual comparison with reference-genome tools

For reference-genome analysis, the closest available comparator is SegMantX (Hanke and Dagan, 2025), which detects diverged segmental duplications using a k-mer chaining approach. As described in the accompanying publication, SegMantX is designed for analysing ancient duplications retained by purifying selection on bacterial plasmids and reports duplication coordinates with an emphasis on chaining diverged matches. It does not classify mechanisms or measure junction microhomology. Tandem Module 1 is designed for the complementary question to classify mechanisms and reports HR signature, microhomology length, and confidence tier per duplication. The two tools therefore occupy different points in the duplication-age vs mechanism-resolution space rather than competing on the same task. We did not run an empirical head-to-head in this work, both because the published descriptions of the tools’ design and intended use already establish their distinct scopes, and because Tandem accepts SegMantX (and other) coordinates via --detection-input, allowing users who already have SegMantX-derived calls on their genome to feed those into Tandem’s mechanism classification stage directly. An empirical evaluation would therefore most usefully take the form of a downstream-pipeline integration test rather than a head-to-head, and is left to future work.

## 4. Discussion

One fundamental assessment in understanding bacterial evolution in growing antibiotics resistance, virulence factor or adaptation is assessing the genome mutations. Among the genomic mutations, duplication is often overlooked not only because of their transient and dynamic nature but also currently there are no tools to assess the genome topology, confirm the junction position and systematically quantify the duplication mutants in populations. Tandem fills a methodological gap in bacterial genomics by providing an integrated pipeline for tandem duplication analysis that goes beyond detection to include mechanism classification and population quantification. The tool’s two-feature classification approach, comprising HR signature at outer boundaries and microhomology length at the inner junction, provides the sequence-level observations needed for mechanism inference without over-claiming pathway identity.

The distinction between Module 1’s two-copy classifier and the single-copy classifier used in Modules 2 and 3 reflects a fundamental biological difference. When two copies are present in the reference, both can be interrogated directly under the R-D-R-D-R bracketing scenarios. When only one copy exists (as in isolate or population analysis against a wild-type reference), the tool measures junction microhomology as end-of-D vs start-of-D and detects HR by searching for an R-D-R structure flanking the single copy in the reference, with adaptive outward windows scaling to duplication size to capture distant flanking repeats such as rRNA operons or IS elements.

Tandem’s HR signature is a statement about the substrate, not about which proteins acted on the junction. An absent HR signature does not equate to RecA inactivity. Tandem’s HR call requires an extended flanking direct repeat (≥ 35 bp at ≥ 92% identity within 5 kb of the boundary), which is the canonical substrate for RecA-mediated homologous recombination as it amplifies, for example, rRNA operons or IS-element pairs. Several alternative mechanisms can produce large tandem duplications without leaving such a footprint. Microhomology-mediated end-joining (MMEJ) operates with 5-25 bp microhomology and is RecA-independent (Chayot, et al., 2010).

Replication template switching at stalled forks can produce structural variants spanning hundreds of kilobases with little or no junction homology (Carvalho and Lupski, 2016; Smith, et al., 2007), and may itself involve RecA acting transiently on the displaced strand without leaving a homology footprint at the final junction (Lovett, 2017). Ku/LigD-type non-homologous end-joining, where present, produces near-blunt junctions (microhomology of 0-4 bp). Tandem reports the sequence-level observations (flanking-repeat presence and junction microhomology) that distinguish among these classes, and in the *P. fluorescens* SBW25 populations the cluster of large duplications (285-767 kb) with both no HR signature and microhomology of 0-1 bp is most consistent with replication-mediated or NHEJ-type formation rather than with classical homology-driven amplification.

Several limitations should be noted. First, Tandem detects forward-orientation flanking repeats consistent with direct (non-inverted) HR. Inverted-repeat-mediated inversion events are outside the scope of this tool. Second, non-templated insertions at junctions (produced by LigD polymerase in organisms with Ku/LigD NHEJ machinery, approximately 20-30% of bacterial species) cannot be detected by reference-based junction-sequence tools, including both Tandem and breseq. Third, static-genome analysis cannot detect duplications that have reverted. Only time-course data can observe reversion directly. Fourth, the HR complexity filter excludes tandem repeat expansions from replication slippage (which is mechanistically distinct from HR), but may also reject HR-mediated by simple repeats such as REP elements. This filter can be disabled with *--no-hr-complexity-filter*. Fifth, Tandem requires the R-D-R geometry to be satisfied on both sides of the duplicated region. One-sided flanking repeats will not produce an HR call even when one boundary clearly sits in a repeat. Finally, circular genome wraparound is handled for distance metrics but not for HR detection windows at genome edges.

Module 2 is designed to confirm the junction position and classify the mechanism rather than a fully automated de novo structural-variant caller. Tandem’s module 2 has two steps. Tandem avoided automizing module 2 and requires user to input the candidate coordinates for junction confirmations. This keeps the user in the loop at the boundary nomination step, which indeed might be the best solution for complicated duplications or raw reads with insufficient coverage. for example, those duplications at highly repetitive regions may not have a clear breakpoint at the boundary or split-read calling that can be automatized. This design choice trades the advantages of automated de novo discovery for two distinct capabilities for further applications: 1) mechanism classification at every confirmed junction, and 2) seamless propagation of confirmed coordinates into Module 3 for population quantification. Module 2 also accepts coordinates from external callers (*breseq*, SegMantX, BISER) via *--detection-input*, so researchers who prefer an automated discovery upstream can pair Tandem with a de novo caller without losing Tandem’s downstream mechanism and frequency analyses. On the *P. fluorescens* SBW25 isolates, Module 2 confirmed all five biological events while *breseq* detected two as new junctions, and Tandem’s mechanism classifications assigned each event to a specific scenario consistent with the published boundary annotations (Khomarbaghi, et al., 2024). Tandem therefore complements rather than replaces general-purpose SV callers.

The JRR is calibrated for short-read paired-end Illumina sequencing of bacterial populations, where per-base error rates of 0.1-1% make exact-match junction counting both fast and accurate. JRR is not directly applicable to long-read populations (Oxford Nanopore, PacBio HiFi). Their higher error rates require error-tolerant junction matching (Hamming or local alignment) to avoid systematic underestimation. Such error-tolerant matching is a planned extension. Within its short-read scope, the 280-test artificial benchmark spans true frequencies of 0-55% across seven bacterial species and four time points (Figure 3A). JRR is systematically lower than the true frequency at every spiked level (a direct consequence of the conservative read-counting design described in Methods section), but is roughly linearly related to it across the range tested (slope ≈ 0.6 in the 0-30% linear range; saturation compression sets in at very high frequencies). The aggregate mean absolute error of 4.3% therefore represents the expected systematic shift, not random scatter. Tandem’s intended use case is therefore tracking duplication dynamics across time-points and conditions, where the conservative bias is consistent and JRR reflects the dynamic of true frequency. JRR measurement improved largely from absolute allele-frequency estimation, which often fail in short-read sequencing or require long-read sequencing with full junction coverage (and error-tolerant matching).

The population-level JRR quantification provided by Module 3 enables a new class of analyses. By quantifying duplication frequency rather than merely detecting presence/absence, users can track duplication dynamics over time in experimental evolution, assess heteroresistance levels in clinical isolates, and study the population genetics of structural variation in natural bacterial communities. Module 3 is, in principle, applicable to any bacterial population sequencing, including the LTEE time-course data of (Good, et al., 2017). Applications to such datasets to characterise duplication dynamics across long-term experimental evolution are an active direction of follow-up work.

Several extensions are priorities for future versions. Empirical validation of the *--coverage-only* workflow on pooled population data is a planned extension. The design admits this use case *(--coverage-only* is sample-type agnostic), but in the present work the artificial population benchmarks were used to validate Module 3’s quantification rather than Module 2’s discovery on pooled samples. Sequencing depth sensitivity has similarly not been characterised systematically. A sweep across 10x, 30x, 100x, and 500x coverage with replicate libraries will quantify the Module 3 detection floor in practice. Validation in repetitive regions, including IS elements, rRNA operons, and prophage remnants, is an explicit gap that future benchmarks will address. The HR classifier currently distinguishes four scenarios (s1-s4) of forward-orientation tandem geometry. Expanding the classifier to capture inverted, nested, partial, and IS-mediated composite duplications would broaden applicability. Finally, replacing the deterministic HR thresholds with a calibrated probabilistic scoring framework, which would combine junction read support, flanking repeat alignment quality, microhomology length, and complexity-filter status into a per-call confidence estimate with reported false-discovery rate, would strengthen downstream meta-analyses and statistical defensibility.

## Acknowledgements

The concept of using Junction Read Ratio (JRR) was discussed and developed from WYN’s PhD work with PhD supervisor Dr. Jenna Gallie at the Max Planck Institute for Evolutionary Biology.

## AI/LLM disclosure

Per ISCB policy on the use of large language models (LLM), the author discloses that an artificial intelligence(AI) assistant (Anthropic Claude and ChatGPT) was used during this work to assist with code review and improve the user interface of Tandem and Grammarly was used for grammar checking and corrections. All scientific reasoning, validation choices, biological interpretation, benchmark design, and final wording were determined by the author. The author takes full responsibility for the accuracy of all content, including code, figures, and text.

## Author Contributions

WYN: Conceptualization; Methodology; Software; Validation; Formal analysis; Investigation; Data curation; Writing (original draft, review and editing); Visualization.

EStJS: Writing (review and editing); Supervision; Project administration.

## Data Availability

Tandem source code, documentation, and example data are freely available at https://github.com/yuingan/tandem under the MIT License (release v0.3.0; archived at the corresponding tag). The *Pseudomonas fluorescens* SBW25 experimental evolution sequencing data analysed in this study are publicly available from the NCBI Sequence Read Archive: isolate sequencing under accessions SRR17278212-SRR17278216 (Khomarbaghi, et al., 2024); day-28 population sequencing under SRR26663277, SRR26663278, and SRR26663279. The reference genome of SBW25 is GenBank accession NC_012660.1. Reference genomes for the seven-species artificial benchmark are listed in Supplementary Table S1. Scripts to reproduce all benchmarks, raw data tables underlying every figure, and the artificial population generator are included in the GitHub repository. All supplementary data are available on Zenodo: 10.5281/zenodo.20345804.

## References

Anderson, R.P. and Roth, J.R. Tandem genetic duplications in Salmonella typhimurium: amplification of the histidine operon. J Mol Biol 1978;126(1):53–71.

Andersson, D.I. and Hughes, D. Gene amplification and adaptive evolution in bacteria. Annu Rev Genet 2009;43:167–195.

Ayan, G.B., Park, H.J. and Gallie, J. The birth of a bacterial tRNA gene by large-scale, tandem duplication events. Elife 2020;9.

Bell, J.C. and Kowalczykowski, S.C. RecA: Regulation and Mechanism of a Molecular Search Engine: (Trends in Biochemical Sciences, June 2016, Vol. 41, No. 6, 491--507). Trends Biochem Sci 2016;41(7):646.

Carvalho, C.M. and Lupski, J.R. Mechanisms underlying structural variant formation in genomic disorders. Nat Rev Genet 2016;17(4):224–238.

Chayot, R., et al. An end-joining repair mechanism in Escherichia coli. Proc Natl Acad Sci U S A 2010;107(5):2141–2146.

Chen, X., et al. Manta: rapid detection of structural variants and indels for germline and cancer sequencing applications. Bioinformatics 2016;32(8):1220-1222.

Cox, M.M. Regulation of bacterial RecA protein function. Crit Rev Biochem Mol Biol 2007;42(1):41–63.

Deatherage, D.E. and Barrick, J.E. Identification of mutations in laboratory-evolved microbes from next-generation sequencing data using breseq. Methods Mol Biol 2014;1151:165–188.

Good, B.H., et al. The dynamics of molecular evolution over 60,000 generations. Nature 2017;551(7678):45–50.

Hanke, D.M. and Dagan, T. SegMantX: A Novel Tool for Detecting DNA Duplications Uncovers Prevalent Duplications in Plasmids. Mol Biol Evol 2025;42(10).

Iseric, H., et al. Fast characterization of segmental duplication structure in multiple genome assemblies. Algorithms Mol Biol 2022;17(1):4.

Khomarbaghi, Z., et al. Large-scale duplication events underpin population-level flexibility in tRNA gene copy number in Pseudomonas fluorescens SBW25. Nucleic Acids Res 2024;52(5):2446–2462.

Layer, R.M., et al. LUMPY: a probabilistic framework for structural variant discovery. Genome Biol 2014;15(6):R84.

Li, H. Minimap2: pairwise alignment for nucleotide sequences. Bioinformatics 2018;34(18):3094–3100.

Li, H., et al. The Sequence Alignment/Map format and SAMtools. Bioinformatics 2009;25(16):2078–2079.

Lovett, S.T. Template-switching during replication fork repair in bacteria. DNA Repair (Amst) 2017;56:118–128.

Maddamsetti, R., et al. Duplicated antibiotic resistance genes reveal ongoing selection and horizontal gene transfer in bacteria. Nat Commun 2024;15(1):1449.

Marcais, G., et al. MUMmer4: A fast and versatile genome alignment system. PLoS Comput Biol 2018;14(1):e1005944.

Rausch, T., et al. DELLY: structural variant discovery by integrated paired-end and split-read analysis. Bioinformatics 2012;28(18):i333–i339.

Sandegren, L. and Andersson, D.I. Bacterial gene amplification: implications for the evolution of antibiotic resistance. Nat Rev Microbiol 2009;7(8):578–588.

Smith, C.E., Llorente, B. and Symington, L.S. Template switching during break-induced replication. Nature 2007;447(7140):102–105.

