## Supplementary for "Tandem: a bioinformatics tool for detection, mechanism classification, and population quantification of bacterial tandem gene duplications"

Wing Yui Ngan

Department of Pharmacology, University of Cambridge, Cambridge, UK

This document contains supplementary tables (S1–S4) and supporting reference materials accompanying the main manuscript.

Raw data files for all benchmarks and the Tandem v0.3.0 source code are available at: https://github.com/Yuingan/Tandem

### Table S1. Complete Module 1 output across eight reference genomes

**Table S1.** Per-duplication output from Tandem Module 1 (v0.3.0, default settings) across all eight reference genomes used in this study. Each row corresponds to one tandem duplication detected by NUCmer self-alignment and classified by the single-copy classifier. Columns: species (with strain identifier); seq_id (reference contig); copy1_start, copy1_end, copy2_start, copy2_end (1-based inclusive boundaries of each duplicate copy); size (duplication length, bp); identity (percent identity of NUCmer alignment); orientation (always "direct" for tandem); distance (gap between copies, bp); is_hr_signature (True if a homologous-recombination signature was detected); hr_match_len (length of the flanking direct repeat, if HR-positive); hr_identity (identity of the flanking repeat alignment); hr_scenario (which of the four single-copy scenarios produced the call); microhomology_bp (length of microhomology at the inner junction); microhomology_seq (sequence of the microhomology); mechanism_confidence (low / moderate / high tier based on duplication size).

| species | size_bp | identity_pct | is_hr | hr_len_bp | hr_id_pct | mh_bp | confidence |
|---|---|---|---|---|---|---|---|
| A. baumannii ATCC17978 | 5694 | 99.9 | False | - | - | 0 | high |
| A. baumannii ATCC17978 | 628 | 82.2 | False | - | - | 4 | high |
| A. baumannii ATCC17978 | 563 | 91.8 | False | - | - | 0 | high |
| A. baumannii ATCC17978 | 512 | 85.2 | False | - | - | 2 | high |
| A. baumannii ATCC17978 | 473 | 89.5 | True | 47 | 93.6 | 0 | moderate |
| A. baumannii ATCC17978 | 423 | 85.7 | False | - | - | 2 | moderate |
| A. baumannii ATCC17978 | 224 | 84.4 | True | 59 | 100.0 | 0 | low |
| E. coli K12 MG1655 | 1059 | 80.8 | False | - | - | 0 | high |
| E. coli K12 MG1655 | 1035 | 97.5 | False | - | - | 2 | high |
| E. coli K12 MG1655 | 885 | 94.2 | False | - | - | 0 | high |
| E. coli K12 MG1655 | 770 | 100.0 | False | - | - | 0 | high |
| E. coli K12 MG1655 | 768 | 98.8 | False | - | - | 0 | high |
| E. coli K12 MG1655 | 380 | 94.5 | False | - | - | 1 | moderate |
| E. coli K12 MG1655 | 358 | 90.8 | False | - | - | 1 | moderate |
| E. coli K12 MG1655 | 354 | 94.4 | True | 183 | 92.3 | 79 | moderate |
| E. coli K12 MG1655 | 295 | 92.9 | True | 53 | 94.3 | 22 | low |
| E. coli K12 MG1655 | 261 | 99.2 | True | 80 | 100.0 | 0 | low |
| E. coli K12 MG1655 | 254 | 85.5 | True | 80 | 100.0 | 0 | low |
| E. coli K12 MG1655 | 212 | 96.7 | True | 106 | 95.3 | 10 | low |
| E. coli K12 MG1655 | 203 | 96.6 | True | 97 | 100.0 | 4 | low |
| E. coli K12 MG1655 | 203 | 95.6 | True | 97 | 99.0 | 25 | low |
| M. genitalium G37 | 858 | 80.0 | False | - | - | 0 | high |
| M. genitalium G37 | 594 | 99.5 | False | - | - | 1 | high |
| M. genitalium G37 | 547 | 86.3 | False | - | - | 0 | high |
| M. genitalium G37 | 261 | 96.5 | True | 69 | 98.6 | 6 | low |
| M. pneumoniae M129 | 1428 | 97.7 | False | - | - | 0 | high |
| M. pneumoniae M129 | 1331 | 93.3 | False | - | - | 0 | high |
| M. pneumoniae M129 | 1237 | 82.3 | False | - | - | 0 | high |
| M. pneumoniae M129 | 1071 | 87.1 | False | - | - | 1 | high |
| M. pneumoniae M129 | 953 | 89.1 | False | - | - | 0 | high |
| M. pneumoniae M129 | 848 | 97.2 | False | - | - | 0 | high |
| M. pneumoniae M129 | 706 | 91.1 | False | - | - | 0 | high |
| M. pneumoniae M129 | 677 | 85.3 | False | - | - | 0 | high |
| M. pneumoniae M129 | 677 | 89.0 | False | - | - | 0 | high |
| M. pneumoniae M129 | 666 | 86.2 | False | - | - | 0 | high |

*(152 rows total; first 35 shown)*

*Full 152-row table with all columns (including coordinates, sequences, and microhomology sequences) is available as TableS1_module1_duplications.tsv in the supplementary data archive.*

### Table S2. HR sensitivity analysis on SBW25

**Table S2.** Sensitivity of Tandem Module 1's HR classification to parameter changes, evaluated on Pseudomonas fluorescens SBW25. Three settings were compared: strict (≥ 50 bp consecutive match at ≥ 95% identity, narrow windows), default (≥ 35 bp consecutive match at ≥ 92% identity, ≤ 5 kb adaptive flanking window), and permissive (relaxed identity and window thresholds). The strict setting also reduces the overall detection pool (from 29 to 11 duplications) by design - strict thresholds are reserved for very high-confidence calls. Default and permissive yielded near-identical mean duplication size (538 bp) and confidence-tier distribution. Columns: setting (strict / default / permissive); n_total (number of duplications detected); n_hr (HR-positive); hr_rate; n_non_hr; mean_size_bp (mean duplication size); conf_low / conf_moderate / conf_high (confidence tier counts).

| setting | n_total | n_hr | hr_rate | n_non_hr | mean_size_bp | conf_low | conf_moderate | conf_high |
|---|---|---|---|---|---|---|---|---|
| strict | 11 | 0 | 0.0000 | 11 | 931 | 0 | 0 | 11 |
| default | 29 | 17 | 0.5862 | 12 | 538 | 12 | 6 | 11 |
| permissive | 29 | 18 | 0.6207 | 11 | 538 | 12 | 6 | 11 |

### Table S3. Runtime benchmarks for all three Tandem modules

**Table S3.** Wall-clock runtime for every benchmark reported in this work, measured with GNU /usr/bin/time -v on the hardware described in §2.7 (AMD Threadripper PRO 7965WX, 44 threads used, 256 GB RAM, Ubuntu 24.04.3 LTS under WSL2). Module 1 scales linearly with genome size and completes all eight reference genomes in under 40 seconds total. Module 2 on SBW25 isolates is comparable to breseq (Tandem mean 45.4 min vs breseq mean 40.2 min per isolate); the small overhead reflects Tandem's additional mechanism classification at every confirmed junction. Module 3 on SBW25 day-28 populations completes in 2.5–3.2 min per population. The full Module 3 artificial benchmark (7 species × 4 timepoints × 10 mutants = 280 tests) completes in approximately 48 minutes total wall time.

| module | test | species_or_isolate | wall_time_s | wall_time_human | notes |
|---|---|---|---|---|---|
| Module 1 | Reference detection | M. genitalium G37 | 2.3 | 2.3 s | 8 reference genomes total |
| Module 1 | Reference detection | M. pneumoniae M129 | 1.8 | 1.8 s | |
| Module 1 | Reference detection | A. baumannii ATCC17978 | 2.9 | 2.9 s | |
| Module 1 | Reference detection | E. coli K12 MG1655 | 3.5 | 3.5 s | |
| Module 1 | Reference detection | M. tuberculosis H37Rv | 5.9 | 5.9 s | |
| Module 1 | Reference detection | P. aeruginosa PAO1 | 5.1 | 5.1 s | |
| Module 1 | Reference detection | P. fluorescens SBW25 | 9.9 | 9.9 s | |
| Module 1 | Reference detection | S. coelicolor A3(2) | 5.7 | 5.7 s | |
| Module 1 | Reference detection total (8 genomes) | - | 37.1 | 37.1 s | |
| Module 2 | SBW25 isolate junction confirmation | SBW25 M1 | 3467.6 | 57.8 min | |
| Module 2 | SBW25 isolate junction confirmation | SBW25 M2 | 2752.8 | 45.9 min | |
| Module 2 | SBW25 isolate junction confirmation | SBW25 M3 | 1861.6 | 31.0 min | |
| Module 2 | SBW25 isolate junction confirmation | SBW25 M4 | 2833.1 | 47.2 min | |
| Module 2 | SBW25 isolate junction confirmation | SBW25 M5 | 2721.5 | 45.4 min | |
| Module 2 | SBW25 total (5 isolates) | - | 13636.6 | 3.79 h | |
| breseq | SBW25 isolate (comparison) | SBW25 M1 | 2321 | 38.7 min | |
| breseq | SBW25 isolate (comparison) | SBW25 M2 | 2430 | 40.5 min | |
| breseq | SBW25 isolate (comparison) | SBW25 M3 | 2307 | 38.5 min | |
| breseq | SBW25 isolate (comparison) | SBW25 M4 | 2458 | 41.0 min | |
| breseq | SBW25 isolate (comparison) | SBW25 M5 | 2544 | 42.4 min | |
| breseq | SBW25 total (5 isolates) | - | 12060 | 3.35 h | |
| Module 3 | SBW25 day-28 population | SBW25 M1 day28 | 194.5 | 3.2 min | |
| Module 3 | SBW25 day-28 population | SBW25 M2 day28 | 152.4 | 2.5 min | |
| Module 3 | SBW25 day-28 population | SBW25 M3 day28 | 155.1 | 2.6 min | |
| Module 3 | SBW25 total (3 populations) | - | 502.0 | 8.4 min | |
| Module 3 | Artificial benchmark (mean of 4 timepoints) | A. baumannii ATCC17978 | 98.1 | 98.1 s | 40 tests per species |
| Module 3 | Artificial benchmark (mean of 4 timepoints) | E. coli K12 MG1655 | 110.2 | 110.2 s | |
| Module 3 | Artificial benchmark (mean of 4 timepoints) | M. genitalium G37 | 15.7 | 15.7 s | |
| Module 3 | Artificial benchmark (mean of 4 timepoints) | M. pneumoniae M129 | 22.1 | 22.1 s | |
| Module 3 | Artificial benchmark (mean of 4 timepoints) | M. tuberculosis H37Rv | 107.3 | 107.3 s | |

| module | test | species_or_isolate | wall_time_s | wall_time_human | notes |
|---|---|---|---|---|---|
| Module 3 | Artificial benchmark (mean of 4 timepoints) | P. aeruginosa PAO1 | 150.9 | 150.9 s | |
| Module 3 | Artificial benchmark (mean of 4 timepoints) | S. coelicolor A3(2) | 217.3 | 217.3 s | |
| Module 3 | Artificial total (28 runs, 7 species × 4 days) | - | 2887 | 48.1 min | |

**Notes on the table:**

• Module 1 was timed with v0.3.0 default settings on each reference genome's main chromosome (plasmids excluded).

• Module 2 SBW25 isolate runs include both Step 1 (coverage exploration via --coverage-only) and Step 2 (junction confirmation with -iso) summed.

• Module 3 SBW25 day-28 populations are the M1, M2, M3 lineages from Khomarbaghi et al. 2024 (NAR), aligned to wild-type SBW25 (NC_012660.1).

• Module 3 artificial-benchmark per-species times are means across the four day-points (day_10, day_20, day_30, day_40). Day_40 is consistently ~50% longer than days 10–30 because it has the highest mutant frequencies (more spanning reads to count and refine).

• breseq was run with v0.39.0 default parameters on the same input read pairs as Tandem Module 2.

#### Table S4. Detailed comparison of Tandem Module 2 vs breseq on SBW25

**Table S4.** Per-junction-variant comparison of Tandem Module 2 (v0.3.0, default settings) and breseq v0.39.0 (default settings) on the five SBW25 evolved isolates (M1–M5) from Khomarbaghi et al. 2024. Each row represents one Tandem-confirmed junction variant. Columns: sample (M1–M5); source (always "tandem" - this is the Tandem call); dup_start, dup_end (1-based inclusive coordinates of the duplicated region); dup_size (bp); spanning_reads (Tandem's exact-match junction read count); is_hr_signature (HR call); microhomology_bp; breseq_status (one of: LD = breseq detected the duplication as elevated read depth; JC_only = breseq detected a junction call near the boundary but did not classify it as a duplication; absent = breseq did not detect this region); breseq_coords (matching breseq coordinates if any).

*Note: M2 has 10 rows here (vs 9 variants reported in main Table 2). The 10th row is a singleton no-HR variant (dup_size 1007434, 1 spanning read) representing a different biological boundary; the main-text Table 2 collapses it into the main 9-variant event. M5 similarly has 11 rows here (vs 11 in main Table 2); the singleton is included as M5_E1.*

| sample | source | tandem_dup_start | tandem_dup_end | tandem_dup_size | tandem_spanning_reads | tandem_is_hr | tandem_microhomology_bp | breseq_status | breseq_coords |
|---|---|---|---|---|---|---|---|---|---|
| M1 | tandem | 1848645 | 2864810 | 1016166 | 21 | False | 3 | JC_only | 1848645/2864810 |
| M2 | tandem | 1852939 | 2865277 | 1012339 | 14 | True | 0 | absent | |
| M2 | tandem | 1852904 | 2865242 | 1012339 | 4 | True | 0 | absent | |
| M2 | tandem | 1852926 | 2865264 | 1012339 | 3 | True | 0 | absent | |
| M2 | tandem | 1852915 | 2865253 | 1012339 | 1 | True | 1 | absent | |
| M2 | tandem | 1852934 | 2865272 | 1012339 | 1 | True | 0 | absent | |
| M2 | tandem | 1852277 | 2865155 | 1012879 | 1 | True | 1 | absent | |
| M2 | tandem | 1852936 | 2865274 | 1012339 | 1 | True | 0 | absent | |
| M2 | tandem | 1849496 | 2856929 | 1007434 | 1 | False | 1 | absent | |
| M2 | tandem | 1852435 | 2865154 | 1012720 | 1 | True | 2 | absent | |
| M2 | tandem | 1852900 | 2865238 | 1012339 | 1 | True | 3 | absent | |
| M3 | tandem | 2296314 | 2786206 | 489893 | 16 | False | 0 | JC_only | 2296498/2786206 |
| M3 | tandem | 2296315 | 2786207 | 489893 | 15 | False | 0 | JC_only | 2296498/2786206 |
| M3 | tandem | 2296223 | 2786988 | 490766 | 3 | True | 0 | absent | |
| M3 | tandem | 2296602 | 2786815 | 490214 | 2 | True | 0 | absent | |
| M3 | tandem | 2296602 | 2786999 | 490398 | 2 | True | 0 | absent | |
| M3 | tandem | 2296332 | 2786912 | 490581 | 1 | True | 2 | absent | |
| M3 | tandem | 2296347 | 2786760 | 490414 | 1 | True | 0 | absent | |
| M3 | tandem | 2296420 | 2787001 | 490582 | 1 | True | 0 | absent | |
| M3 | tandem | 2296438 | 2786835 | 490398 | 1 | True | 0 | absent | |
| M3 | tandem | 2296468 | 2786865 | 490398 | 1 | True | 0 | absent | |
| M3 | tandem | 2296403 | 2786984 | 490582 | 1 | True | 0 | absent | |
| M3 | tandem | 2296265 | 2786846 | 490582 | 1 | True | 0 | absent | |
| M3 | tandem | 2296245 | 2786826 | 490582 | 1 | True | 0 | absent | |
| M3 | tandem | 2296600 | 2786813 | 490214 | 1 | True | 2 | absent | |

| sample | source | tandem_dup_start | tandem_dup_end | tandem_dup_size | tandem_spanning_reads | tandem_is_hr | tandem_microhomology_bp | breseq_status | breseq_coords |
|---|---|---|---|---|---|---|---|---|---|
| M4 | tandem | 1852991 | 2865329 | 1012339 | 1 | True | 1 | absent | |
| M4 | tandem | 1852949 | 2865557 | 1012609 | 1 | True | 1 | absent | |
| M5 | tandem | 1854447 | 2866785 | 1012339 | 10 | True | 0 | absent | |
| M5 | tandem | 1854398 | 2866736 | 1012339 | 3 | True | 0 | absent | |
| M5 | tandem | 1854438 | 2866776 | 1012339 | 2 | True | 1 | absent | |
| M5 | tandem | 1854409 | 2866747 | 1012339 | 1 | True | 0 | absent | |
| M5 | tandem | 1854402 | 2866740 | 1012339 | 1 | True | 1 | absent | |
| M5 | tandem | 1854406 | 2866744 | 1012339 | 1 | True | 0 | absent | |
| M5 | tandem | 1854230 | 2866568 | 1012339 | 1 | True | 0 | absent | |
| M5 | tandem | 1852435 | 2865154 | 1012720 | 1 | True | 2 | absent | |
| M5 | tandem | 1854272 | 2866610 | 1012339 | 1 | True | 1 | absent | |
| M5 | tandem | 1854332 | 2866670 | 1012339 | 1 | True | 0 | absent | |
| M5 | tandem | 1854597 | 2866935 | 1012339 | 1 | True | 0 | absent | |

#### Script S1. Benchmark scripts

**Script S1.** Bash scripts that reproduce all benchmarks reported in this manuscript. These are included in the GitHub repository (https://github.com/yuingan/tandem) and are not embedded here. Files:

- run_v0.3_sbw25.sh: Module 2 + breseq comparison on SBW25 M1–M5 isolates
- run_artificial_benchmark.sh: Module 3 artificial population benchmark on seven bacterial species × four time points × ten mutants per species
- run_module1_benchmark.sh: Module 1 detection on eight reference genomes (default and HR sensitivity comparisons)

Each script logs all Tandem output and time -v measurements into a logs/ subdirectory, and writes summary TSVs that feed into Tables 1, 2, 3, S1, S2, S3, and S4 as well as Figures 3 and 4.
